# Microglia-derived IL-18 remodels hippocampal plasticity to constrain traumatic fear memory

**DOI:** 10.64898/2026.05.04.721266

**Authors:** Xiya Fang, Yining Wang, Yezhuang Shen, Guanbo Xie, Ran Liu, Hong Cai, Qiuying Han, Xin Xu, Kai Wang, Libing Yin, Jinwei Di, Tao Zhou, Ailing Li, Xiao Han, Weihua Li, Teng Li

## Abstract

Post-traumatic stress disorder (PTSD) involves complex neuroimmune-synaptic crosstalk in the hippocampus. Here we show that interleukin-18 (IL-18), an extensively studied pro-inflammatory cytokine, serves a protective role against traumatic fear memory through a microglia-to-neuron signaling axis. Traumatic stress induces sustained upregulation of IL-18 in the hippocampus. Exogenous IL-18 administration attenuates fear memory, whereas blockade of IL-18 signaling exacerbates it. Mechanistically, microglial-derived IL-18 acts on neuronal IL-18R1 to restore stress-impaired synaptic plasticity and reduce perineuronal net density, thereby facilitating structural synaptic remodeling. In addition, IL-18 modulates the synaptic organization of fear memory-encoding engram cells within hippocampal ensembles. Together, these findings redefine IL-18 as a homeostatic regulator of post-trauma hippocampal synaptic function and identify the hippocampal IL-18 pathway as a potential therapeutic target for PTSD.

## Introduction

Post-traumatic stress disorder (PTSD) is a debilitating psychiatric condition triggered by severe traumatic events, characterized by intrusive re-experiencing of traumatic memories, persistent hyperarousal, and pronounced avoidance behaviors, imposing substantial disease burden on both individuals and society^1,2^. The societal impact of this disorder is profound and multidimensional: at the individual level, patients frequently suffer from severe cognitive impairment, dysregulated emotional control, and deteriorated social functioning, with approximately half of cases progressing to chronic disability due to protracted disease courses; at the public health level, PTSD exhibits extraordinarily high comorbidity rates with major depression, generalized anxiety disorder, and substance use disorders, significantly compounding disease complexity and treatment challenges^3,4^. Currently, the pathological mechanisms underlying PTSD remain incompletely understood, and clinically effective disease-specific therapies are lacking^5,6^.

Neuroimmune dysregulation has emerged as a fundamental contributor to PTSD pathogenesis, challenging the long-held view that this disorder arises primarily from intrinsic neuronal dysfunction^4,7^. Accumulating evidence demonstrates that traumatic stress activates microglia and astrocytes, initiating multi-layered immune cascades that encompass NLRP3 inflammasome assembly, release of pro-inflammatory cytokines (e.g., IL-1β, IL-6, TNF-α)^8–10^, upregulation of chemokines (e.g., CCL2, CXCL1), and components of the classical complement cascade (C1q, C3), which collectively reshape the post-traumatic neuroimmune landscape and drive aberrant microglia-mediated synaptic remodeling^11–13^. These immune mediators not only drive neuroinflammation but also directly modulate synaptic plasticity and neuronal function, impairing hippocampus-dependent memory extinction, potentiating amygdala-mediated fear responses, and forming vicious cycles with hypothalamic-pituitary-adrenal (HPA) axis dysfunction, collectively establishing the pathological foundation for consolidation and maintenance of maladaptive fear memories^14–16^. However, the role of neuroimmune signaling is not simply linear or uniformly deleterious; rather, its effects exhibit high context-dependence and complexity, suggesting the existence of homeostatic regulatory mechanisms that remain insufficiently appreciated^17^.

Among stress-responsive cytokines, interleukin-18 (IL-18) occupies a paradoxical position^18^. Traditionally, IL-18 has been regarded as a pro-inflammatory cytokine^19^. Clinical observations reveal elevated serum IL-18 levels in PTSD patients, and animal studies demonstrate stress-induced upregulation of IL-18 expression in brain regions such as the amygdala, collectively supporting its potential involvement in disease progression^20^. However, contradictory evidence continues to emerge: IL-18-deficient mice paradoxically exhibit exacerbated neuroinflammation, HPA axis hyperactivation, and anxiety-like behaviors following acute stress^21,22^. Such discrepancies likely reflect heterogeneity in species, stress paradigms, brain regions, and measurement timing across studies, rather than mechanistic conflict. Even so, the temporal trajectory and cellular source of hippocampal IL-18 after traumatic stress, its effect on synaptic plasticity, and whether it acts on fear memory-encoding neurons all remain undefined.

At the cellular level, the hippocampus is acutely vulnerable to traumatic stress^23^. Stress disrupts glutamatergic transmission and long-term potentiation, in part through loss of presynaptic vesicular glutamate transporters and postsynaptic scaffolding proteins that maintain synaptic integrity^9,24^. In parallel, stress remodels the extracellular matrix, particularly perineuronal nets (PNNs), condensed proteoglycan assemblies that stabilize mature synaptic contacts and restrict plasticity. Stress-induced upregulation of PNNs is thought to consolidate pathological fear memories and impair fear extinction, the deficit that defines PTSD^25^. However, whether and how immune signaling in the stressed hippocampus coordinates glutamatergic synaptic remodeling and PNN regulation, and whether such coordination contributes to the attenuation of traumatic fear memory, remains unknown.

Fear memories can be encoded by sparse hippocampal engram ensembles whose synaptic organization determines memory specificity^26,27^. Engrams, defined as sparse neuronal ensembles activated during encoding and reactivated during retrieval, represent the physical trace of fear memories^28^. During fear conditioning, hippocampal neurons with higher intrinsic excitability are preferentially recruited into engram ensembles, exhibiting enhanced synaptic connectivity^29,30^. In PTSD, excessive memory consolidation, impaired extinction, and aberrant engram reactivation sustain pathological fear responses^31,32^.

Emerging evidence suggests that the precise tuning of synaptic weights within engram circuits, rather than their overall strengthening, determines whether fear memories become generalized or context-specific^33,34^. Whether IL-18-mediated neuroimmune signaling can shape the synaptic organization of hippocampal engram ensembles during fear memory consolidation remains entirely unknown.

Here, we demonstrate that hippocampal IL-18 functions as a protective factor against PTSD-related fear memory through a microglia-to-neuron signaling axis. We find that traumatic stress triggers sustained IL-18 upregulation in the hippocampus, and that exogenous IL-18 attenuates fear memory while blockade of IL-18 signaling exacerbates it. At the synaptic level, IL-18 restores stress-impaired synaptophysin expression and enhances functional excitatory synaptic connectivity marked by VGLUT1–Homer1 co-localization. IL-18 also reduces PNNs density in the hippocampus, facilitating structural synaptic remodeling. At the engram level, IL-18 treatment decreases the number of trauma-activated engram cells in the hippocampus while simultaneously augmenting VGLUT1 expression within these ensembles. Together, these findings redefine IL-18 as a homeostatic neuroimmune regulator that reshapes synaptic architecture and engram synaptic organization to constrain pathological fear memory, and identify the hippocampal IL-18 pathway as a potential therapeutic target for PTSD.

## Results

### Elevation of Hippocampal IL-18 During the Development of PTSD-like Behavior in Mice

We first established a mouse PTSD model using single prolonged stress combined with foot shock (SPS&S), which recapitulates the clinical features of PTSD patients^35,36^. This paradigm comprises four consecutive stressors (one-trial 2-hour restraint, 15-minute forced swim, ether anesthesia, and 5-minute exposure to a context followed by a 10-second, 0.8 mA foot shock with constant current), followed by behavioral assessments for contextual fear, depression, and anxiety (Fig. 1a and Supplementary Fig.1a). SPS&S-treated mice exhibited significantly enhanced fear memory and depression- and anxiety-like behaviors at both day 8 and day 15 post-stress, recapitulating core behavioral features observed in clinical PTSD and confirming the utility of this paradigm for modeling post-traumatic stress responses (Fig. 1a-c and Supplementary Fig.1b-g).

**Fig. 1.**
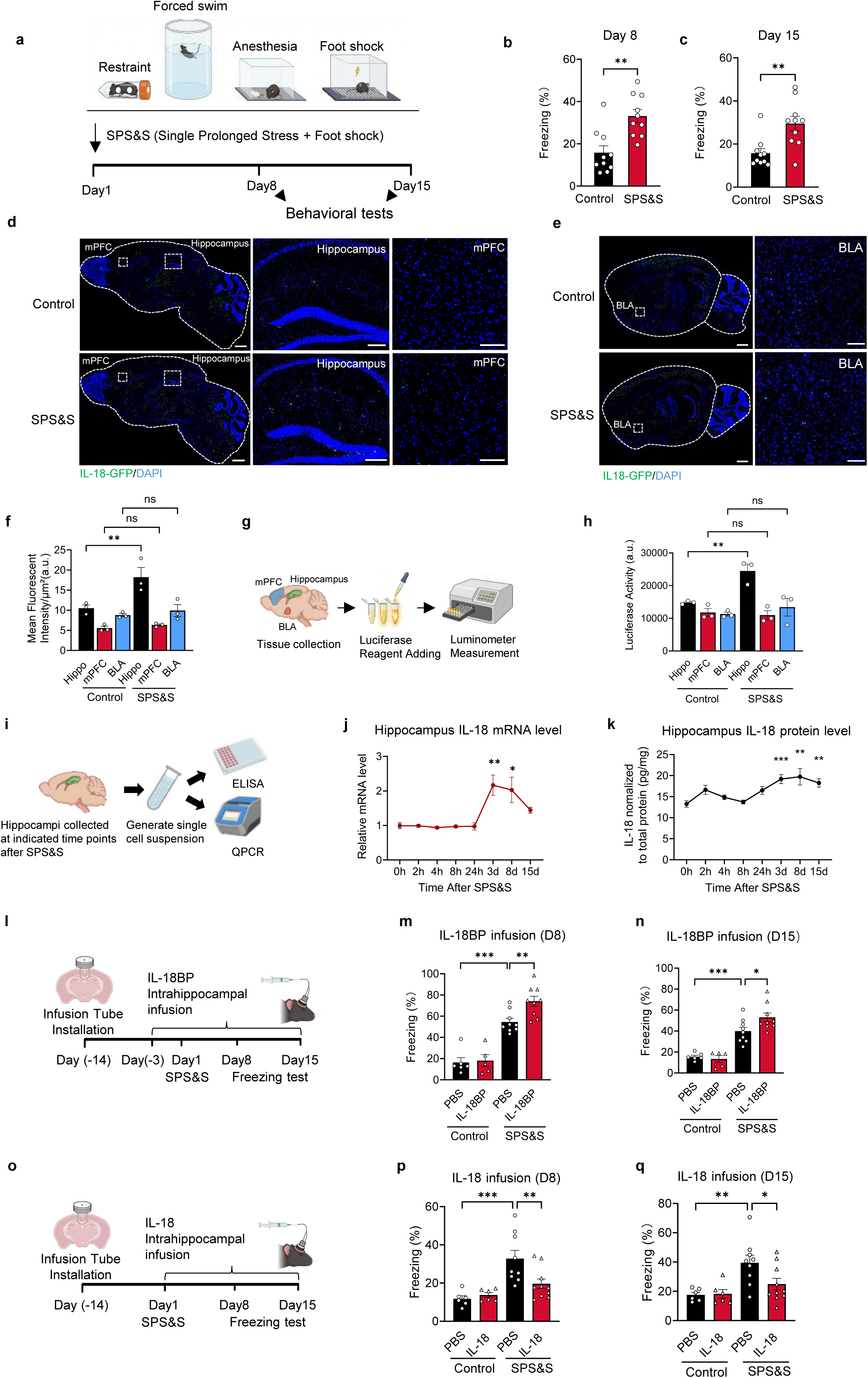
Stress-induced hippocampal IL-18 upregulation constrains traumatic fear memory formation. **a**, Schematic of PTSD model establishment. The SPS&S (Single Prolonged Stress & Shock) procedure, consisting of sequential restraint, forced swim, anesthesia, and foot shock. **b–c**, Contextual fear freezing in SPS&S-exposed and control mice at day 8 (**b**) and day 15 (**c**) post-stress. **P < 0.01, unpaired two-tailed t-test; n = 10 mice per group. **d–e**, Representative images showing IL-18-GFP fluorescence in sagittal brain sections from IL-18-GFP reporter mice at day 3 post-SPS&S. Images show whole-brain overview with magnified insets of the hippocampus and medial prefrontal cortex (mPFC) (**d**), and basolateral amygdala (BLA). IL-18-GFP specks are represented by green dots generated using IMARIS software. (**e**), scale bars,1mm (overview), 200 μm (inset, hippocampus), 100 μm (inset, mPFC and BLA). **f**, Quantification of mean IL-18-GFP fluorescent intensity across hippocampus, mPFC, and BLA in control and SPS&S groups. **P < 0.01, ns, not significant; two-way ANOVA with Bonferroni’s post hoc test; n = 3 mice per group. **g**, Schematic of the IL-18 luciferase reporter assay for quantifying IL-18 bioactivity in dissected brain regions. **h**, Luciferase activity in hippocampus, mPFC, and BLA from control and SPS&S mice. **P < 0.01, ns, not significant; two-way ANOVA with Bonferroni’s post hoc test; n = 3 per group.**i**, Schematic of hippocampal tissue collection and processing for IL-18 mRNA quantification by qPCR and protein quantification by ELISA at multiple post-stress time points. n = 3 mice per group. **j**, Temporal profile of hippocampal IL-18 mRNA levels from 0 to 15 days post-SPS&S. *P < 0.05, **P < 0.01 vs. 0 h; one-way ANOVA with Dunnett’s post hoc test; n = 7-11 per time point. **k**, Temporal profile of hippocampal mature IL-18 protein levels (normalized to total protein) from 0 to 15 days post-SPS&S. **P < 0.01 vs. 0 h; one-way ANOVA with Dunnett’s post hoc test; n = 7-11 per time point. **l**, Experimental timeline for intrahippocampal cannula implantation, SPS&S exposure, and daily PBS or IL-18 binding protein (IL-18BP) infusion, with fear memory assessed at days 8 and 15 post-stress. IL-18BP infusion began on day −3 to allow accumulation of antagonist before stress onset, ensuring complete neutralization of stress-induced IL-18. **m–n**, Contextual fear freezing at day 8 (**m**) and day 15 (**n**) in mice receiving intrahippocampal IL-18BP or PBS following SPS&S. *P < 0.05, **P < 0.01, ***P < 0.001; two-way ANOVA with Bonferroni’s post hoc test; n = 5-9 per group. **o**, Experimental timeline for intrahippocampal cannula implantation, SPS&S exposure, and daily IL18 infusion, with fear memory assessed at days 8 and 15 post-stress. **p–q**, Contextual fear freezing at day 8 (**p**) and day 15 **(q**) in mice receiving intrahippocampal IL18 or PBS following SPS&S. *P < 0.05, **P < 0.01, ***P < 0.001; two-way ANOVA with Bonferroni’s post hoc test; n = 6–10 per group. Complete statistics are provided in Supplementary Table 1.

Given that IL-18 has been implicated in anxiety- and depression-like behavior regulation ^37^, and that various stress paradigms increase IL-18 expression across brain regions, we first examined the spatial dynamics of IL-18 following SPS&S exposure. Using IL18-Luc-2A-EGFP reporter mice (IL-18-GFP reporter mice), whole-brain imaging revealed that SPS&S obviously upregulated IL-18 in the hippocampus, with no significant changes observed in other stress-responsive regions including the medial prefrontal cortex (mPFC) and basolateral amygdala (BLA) (Fig. 1d–f). To further validate this regional specificity, we isolated hippocampal, mPFC, and BLA tissues and quantified luciferase activity ex vivo (Fig. 1g). Consistent with the immunostaining results, luciferase signals were significantly elevated in the hippocampus but not in the mPFC or BLA following SPS&S (Fig. 1h). Together, these findings indicate that SPS&S-induced IL-18 upregulation is preferentially localized to the hippocampus among the three key stress-responsive regions examined.

We next examined the temporal dynamics of hippocampal IL-18 following SPS&S by measuring IL-18 mRNA and mature protein levels at multiple post-stress time points (Fig. 1i). Both the mRNA and mature forms IL-18 protein were significantly elevated at day 3 and remained upregulated through day 8, with mature IL-18 protein persisting up to at least day 15 (Fig. 1j, k). In contrast, hippocampal IL-1β, another NLRP3 inflammasome substrate, peaked earlier at 2 hours post-stress (Supplementary Fig. 1h), revealing a clear temporal dissociation between IL-18 and IL-1β induction despite their shared upstream pathway. Notably, the sustained elevation of hippocampal IL-18 aligns with the time course of PTSD-like symptomatology, suggesting that IL-18 signaling may actively participate in the neurobiological processes underlying PTSD.

### Hippocampal IL-18 Signaling Regulates Traumatic Fear Memory

To directly test whether endogenous hippocampal IL-18 signaling is required for fear memory regulation, we implanted bilateral cannulae targeting the dorsal hippocampus and administered IL-18 binding protein (IL-18BP), a natural IL-18 antagonist, to neutralize endogenous IL-18 prior to SPS&S exposure and daily thereafter, with contextual fear memory assessed at days 8 and 15 post-stress (Fig. 1l). IL-18BP-treated mice displayed markedly enhanced fear freezing compared to vehicle controls at both time points (Fig. 1m, n), demonstrating that disruption of endogenous IL-18 signaling exacerbates pathological fear memory. Neither IL-18BP nor vehicle administration affected overall locomotor activity during the testing period (Supplementary Fig. 1i, j). To further examine the functional consequence of exogenous IL-18 supplementation on traumatic fear memory, mice received intrahippocampal IL-18 or vehicle immediately before SPS&S exposure and daily thereafter (Fig. 1o). IL-18-treated mice exhibited significantly reduced fear freezing compared to PBS controls at days 8 and 15 post-stress (Fig. 1p, q), confirming that exogenous IL-18 attenuates fear memory formation. Locomotor activity remained unaffected by IL-18 administration (Supplementary Fig. 1k, l). Together, these results establish hippocampal IL-18 as a regulator of traumatic fear memory.

### Microglia-Derived IL-18 in the Hippocampus Regulates Fear Memory

It has been reported that IL-18 release in the central nervous system involves coordinated responses from multiple glial cell populations^38,39^. Following restraint stress, microglia serve as the predominant source of IL-18, while astrocytes can also upregulate IL-18 production upon inflammatory stimulation^38^. To identify the primary cellular source of hippocampal IL-18 following SPS&S, we subjected IL-18-GFP reporter mice to stress and performed immunostaining 3 days later. IL-18 expression co-localized predominantly with microglia, and to a lesser extent with astrocytes and neurons (Fig. 2a). To establish the functional relevance of this cellular source, we employed AAV-mediated shRNA to selectively knock down IL-18 in neurons, microglia, or astrocytes prior to SPS&S exposure, and assessed contextual fear memory at days 8 and 15 post-stress (Fig. 2b). Cell-type-specific depletion was confirmed by immunostaining (Supplementary Fig. 2a). Microglia-specific IL-18 knockdown significantly enhanced contextual fear freezing at both days 8 and 15 post-SPS&S, whereas IL-18 depletion in astrocytes or neurons had no discernible effect on fear memory (Fig. 2c, d). These results identify microglia as the functionally relevant source of hippocampal IL-18 in the context of PTSD-like fear memory.

**Fig. 2.**
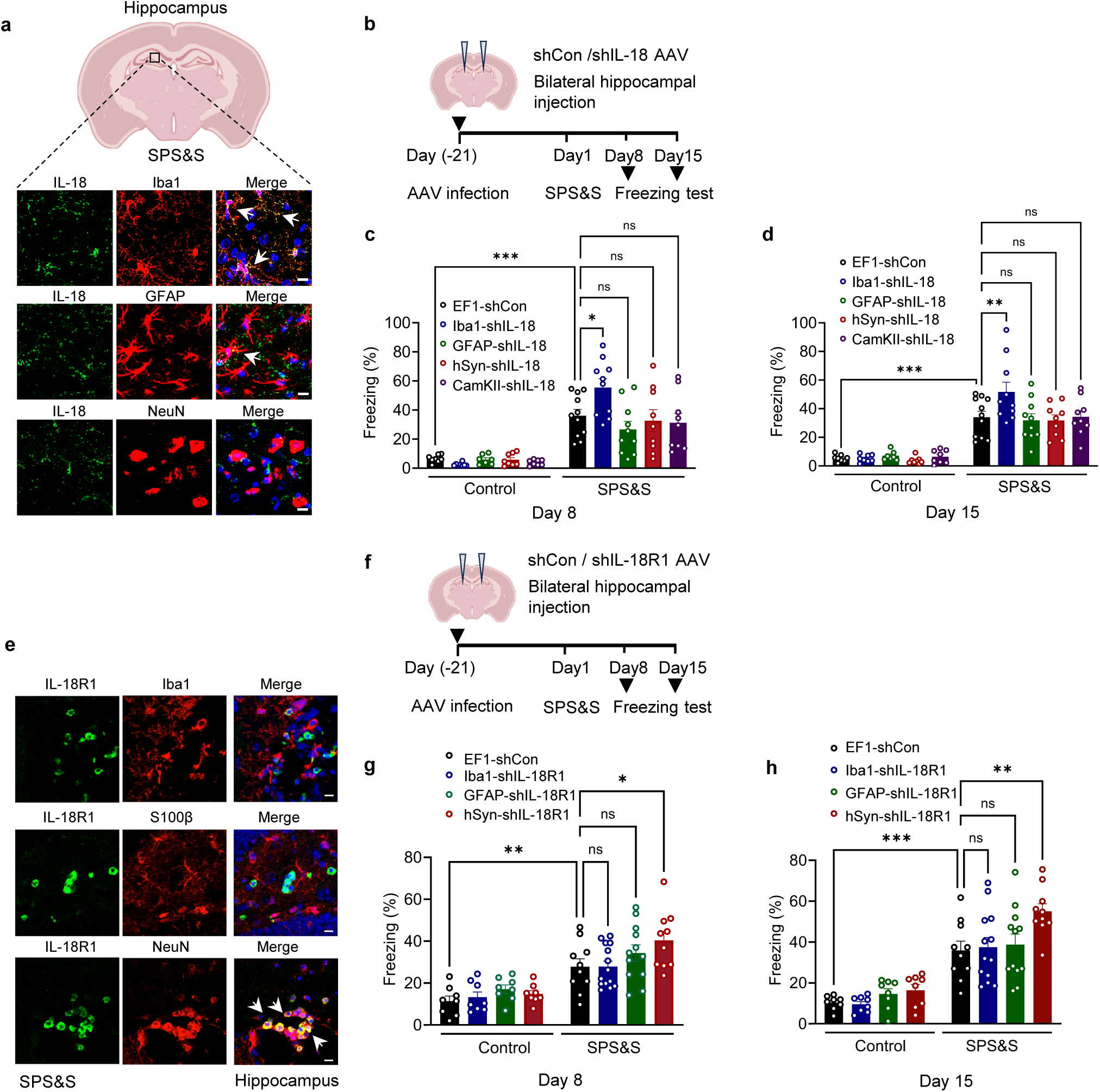
Microglia-derived IL-18 regulates fear memory via neuronal IL-18R1 signaling in the hippocampus. **a,** Representative confocal images of hippocampal sections from IL-18-GFP reporter mice at day 3 post-SPS&S. Scale bars, 10 μm. **b,** Experimental timeline for bilateral hippocampal AAV injection (day −21), SPS&S exposure (day 1), and contextual fear memory assessment (days 8 and 15). **c,** Contextual fear freezing at day 8 in mice receiving cell-type-specific AAV-shRNA targeting IL-18 in microglia (Iba1-shIL18), astrocytes (GFAP-shIL18), neurons (hSyn-shIL18 and CaMKII-shIL18), or scramble control (EF1-shCon), following SPS&S or control conditions. *P < 0.05, **P < 0.01, ***P < 0.001, ns, not significant Two-way ANOVA with post-hoc Bonferroni correction, n=7-11 mice per group. **d,** Same as **c** at day 15. **e,** Representative confocal images of hippocampal sections stained for IL-18R1. Scale bars, 10 μm. **f,** Experimental timeline for bilateral hippocampal AAV injection (day −21), SPS&S exposure (day 1), and contextual fear memory assessment (days 8 and 15) for IL-18R1 knockdown experiments. **g,** Contextual fear freezing at day 8 in mice receiving cell-type-specific AAV-shRNA targeting IL-18R1 in microglia (Iba1-shIL18R1), astrocytes (GFAP-shIL18R1), or neurons (hSyn-shIL18R1), or scramble control (EF1-shCon), following SPS&S or control conditions. *P < 0.05, **P < 0.01, ***P < 0.001, ns, not significant Two-way ANOVA with post-hoc Bonferroni correction, n=8-13 mice per group. **h,** Same as **g** at day 15. Complete statistics are provided in Supplementary Table 1.

### Neuronal IL-18R1 Is Required for Hippocampal IL-18’s Regulation of Fear Memory

Having identified microglia as the cellular source of stress-induced hippocampal IL-18 (Fig. 2a–d), we next asked which cell type receives this signal. Immunostaining revealed that IL-18R1 was expressed predominantly by neurons, with modest expression in microglia and astrocytes (Fig. 2e). To determine which receptor-expressing population is functionally required, we used AAV-mediated shRNA to selectively knock down IL-18R1 in hippocampal neurons, microglia, or astrocytes prior to SPS&S exposure (Fig. 2f). Cell-type-specific knockdown efficiency was validated by immunostaining (Supplementary Fig. 2b). Neuron-specific IL-18R1 depletion significantly enhanced contextual fear responses at days 8 and 15 post-stress, whereas depletion in microglia or astrocytes produced no detectable effect (Fig. 2g, h). Together, these findings complete a microglia-to-neuron signaling axis in which microglia-derived IL-18 engages neuronal IL-18R1 to constrain hippocampal fear memory.

### Neuronal IL-18R1 Signaling Is Required for Maintaining Hippocampal Synaptic Integrity Following Traumatic Stress

Immune system cytokines can modulate synaptic development, plasticity, and cognition in the central nervous system^40,41^. We next examined whether this microglial-neuronal IL-18/IL-18R1 signaling axis underlies changes in hippocampal synaptic plasticity following SPS&S. Synaptophysin was used as a general presynaptic terminal marker, VGLUT1 as an indicator of presynaptic glutamate packaging capacity, and Homer1 as a postsynaptic scaffold protein that anchors metabotropic glutamate receptors at the postsynaptic density and links intracellular signaling to calcium stores^42,43^. SPS&S exposure significantly reduced synaptophysin expression and VGLUT1/Homer1 co-localization puncta, reflecting disruption of glutamatergic synaptic integrity (Fig. 3a-g). AAV-mediated IL-18R1 knockdown in hippocampal neurons further exacerbated these deficits, significantly decreasing the stress-induced decrease in synaptophysin expression (Fig. 3b-d), accompanied by a trend toward decreased VGLUT1/Homer1 co-localization and VGLUT1 expression (Fig. 3f, g), while Homer1 levels remained unaffected (Supplementary Fig. 3a). Together, these findings indicate that neuronal IL-18R1 signaling is required to maintain glutamatergic synaptic integrity under traumatic stress, and its disruption exacerbates stress-induced synaptic impairment.

**Fig. 3.**
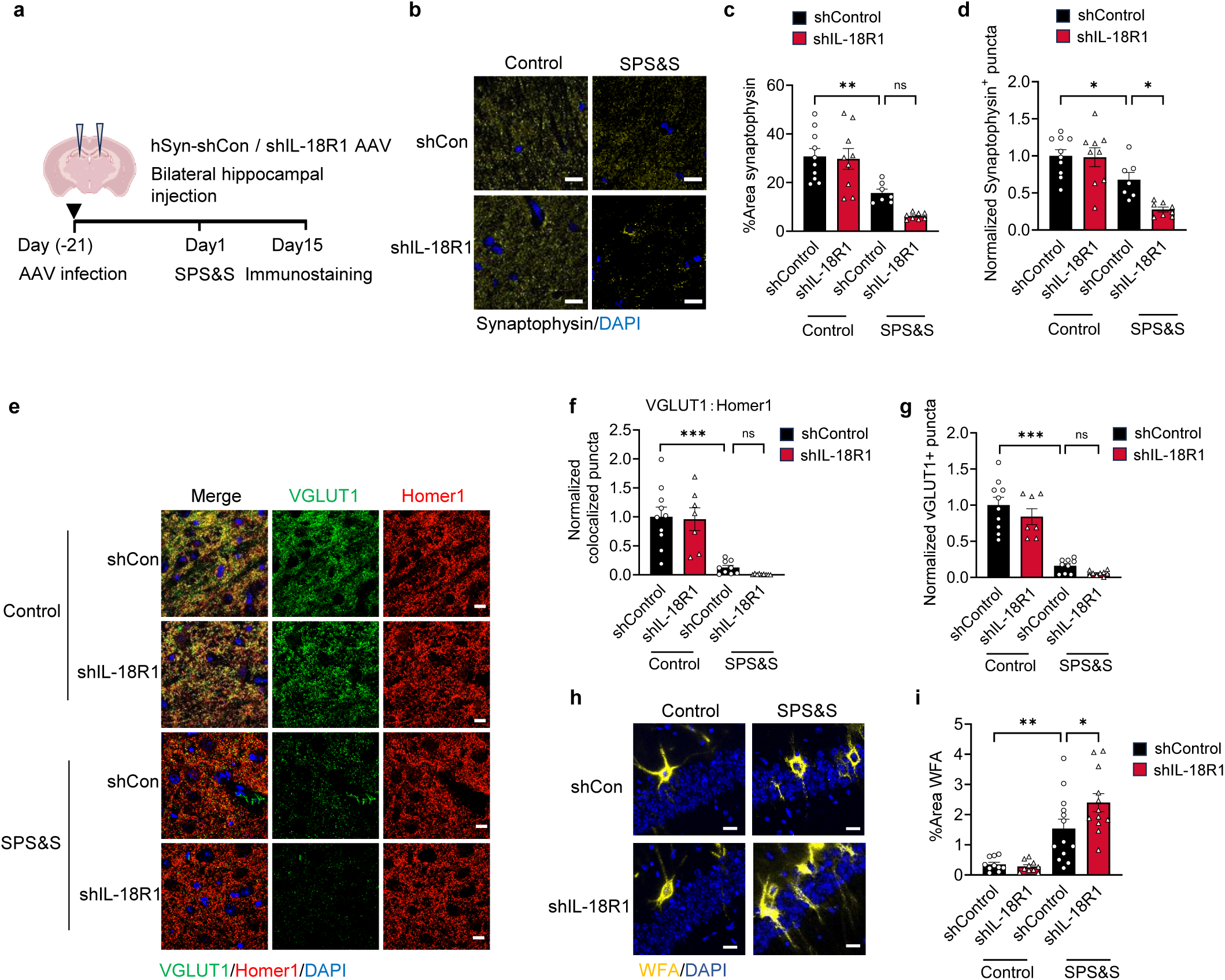
Neuronal IL-18R1 signaling is required for maintaining hippocampal synaptic integrity following traumatic stress. **a,** Experimental timeline for bilateral hippocampal AAV-shRNA injection (day −21), SPS&S exposure (day 1), and tissue collection for immunostaining at day 15. **b,** Representative confocal images of synaptophysin (yellow) and DAPI (blue) immunostaining in hippocampal CA1 from shControl and shIL-18R1 mice under control and SPS&S conditions. Scale bars, 10 μm. **c,** Quantification of percent area synaptophysin staining across groups. *P < 0.05, **P < 0.01, Two-way ANOVA with post-hoc Holm-Šídák, n=7-10 fields of view (FOVs) per group, from 3 mice per group. **d,** Quantification of normalized Synap⁺ puncta density. *P < 0.05, Two-way ANOVA with post-hoc Bonferroni correction, n=7-10 fields of view (FOVs) per group, from 3 mice per group. **e,** Representative confocal images of VGLUT1 (green), Homer1 (red), and DAPI (blue) co-immunostaining in hippocampal CA1 from shControl and shIL-18R1 mice under control and SPS&S conditions. Scale bars, 10 μm. **f,** Quantification of normalized VGLUT1–Homer1 co-localization puncta. ***P < 0.001, ns, not significant; Two-way ANOVA with post-hoc Bonferroni correction, n=7-10 fields of view (FOVs) per group, from 3 mice per group. **g,** Quantification of normalized VGLUT1⁺ puncta density. ***P < 0.001, ns, not significant, Two-way ANOVA with post-hoc Bonferroni correction, n=7-10 fields of view (FOVs) per group, from 3 mice per group. **h,** Representative confocal images of WFA (yellow) and DAPI (blue) staining in hippocampal CA1 from shControl and shIL-18R1 mice under control and SPS&S conditions. Scale bars, 20 μm. **i,** Quantification of percent area WFA staining. *P < 0.05, **P < 0.01; two-way ANOVA with post-hoc Bonferroni correction, n=10-13 fields of view (FOVs) per group, from 3 mice per group. Data are presented as mean ± SEM. Complete statistics are provided in Supplementary Table 1.

Extensive evidence demonstrates that the extracellular matrix (ECM), of which perineuronal nets (PNNs) are a critical component, undergoes dynamic regulation following learning experiences, encoding experience into memory through synaptic remodeling^44,45^. To determine whether neuronal IL-18 signaling contributes to ECM homeostasis following stress, we assessed hippocampal PNN density using Wisteria floribunda agglutinin (WFA) staining and aggrecan immunostaining. Neuronal IL-18R1 knockdown increased both WFA staining intensity and aggrecan expression following SPS&S (Fig. 3h, i and Supplementary Fig. 3b, c), indicating that loss of neuronal IL-18 receptor signaling elevates PNN density and impairs the structural remodeling capacity necessary for adaptive synaptic plasticity.

### Exogenous IL-18 Restores Stress-Impaired Synaptic Plasticity and Extracellular Matrix Remodeling

We next examined whether exogenous IL-18 administration influences stress-impaired synaptic plasticity in the hippocampus. Mice received IL-18 or vehicle immediately before SPS&S exposure and daily thereafter via pre-implanted bilateral cannulae (Fig. 4a). IL-18 treatment significantly attenuated the stress-induced reduction in synaptophysin expression (Fig. 4b–d). At the level of excitatory synaptic connectivity, IL-18 administration significantly rescued both VGLUT1–Homer1 co-localization puncta and VGLUT1 expression following SPS&S (Fig. 4e-g), while Homer1 expression remained unchanged (Supplementary Fig. 3d). Importantly, IL-18 treatment significantly reduced WFA staining intensity and aggrecan expression following SPS&S (Fig. 4h, i and Supplementary Fig. 3e, f), suggesting that IL-18 promotes PNN remodeling to facilitate synaptic structural plasticity. Together, these findings demonstrate that hippocampal IL-18 signaling regulates post-traumatic synaptic plasticity through coordinated modulation of synaptophysin expression, excitatory synaptic connectivity, and perineuronal net remodeling.

**Fig. 4.**
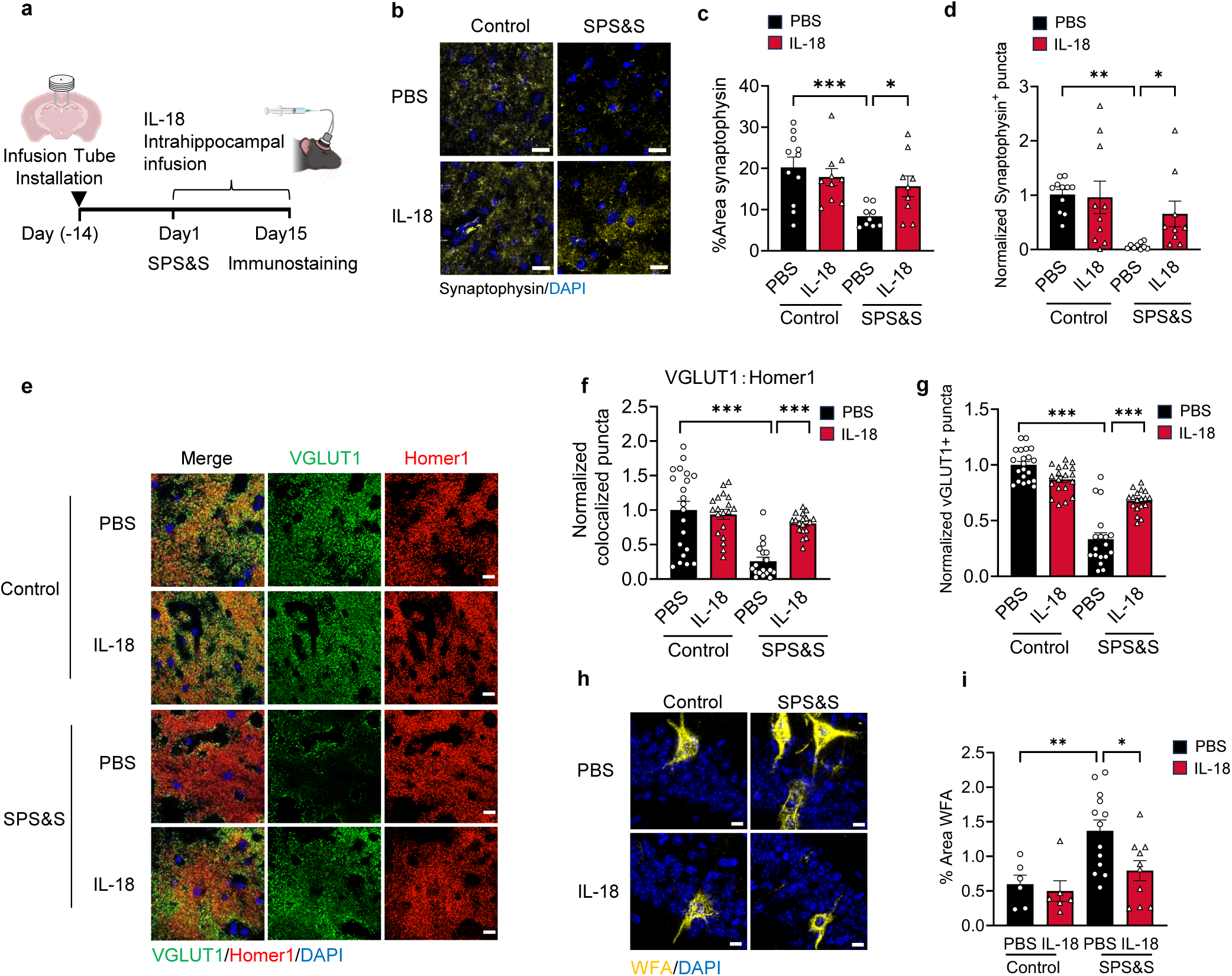
Exogenous IL-18 restores stress-impaired synaptic plasticity and extracellular matrix remodeling. **a,** Experimental timeline for infusion tube installation (day −14), SPS&S exposure (day 1), daily intrahippocampal IL-18 or PBS infusion, and tissue collection for immunostaining at day 15. **b,** Representative confocal images of synaptophysin (yellow) immunostaining in hippocampal CA1 from PBS- and IL-18-treated mice under control and SPS&S conditions. Scale bars, 10 μm. **c,** Quantification of percent area synaptophysin staining across groups. ***P < 0.001, *P < 0.05, Two-way ANOVA with post-hoc Holm-Šídák, n=9-11 fields of view (FOVs) per group, from 3 mice per group. **d,** Quantification of normalized synaptophysin⁺ puncta density. Two-way ANOVA with post-hoc Holm-Šídák, n=9-11 fields of view (FOVs) per group, from 3 mice per group. **e,** Representative confocal images of VGLUT1 (green), Homer1 (red), and DAPI (blue) co-immunostaining in hippocampal CA1 from PBS- and IL-18-treated mice under control and SPS&S conditions. Scale bars, 10 μm. **f,** Quantification of normalized VGLUT1–Homer1 co-localization puncta. ***P < 0.001, Two-way ANOVA with post-hoc Bonferroni correction, n=18-21 fields of view (FOVs) per group, from 3 mice per group. **g,** Quantification of normalized VGLUT1⁺ puncta density. ***P < 0.001, Two-way ANOVA with post-hoc Bonferroni correction, n=18-21 fields of view (FOVs) per group, from 3 mice per group. **h,** Representative confocal images of WFA (yellow) and DAPI (blue) staining in hippocampal CA1 from PBS- and IL-18-treated mice under control and SPS&S conditions. Scale bars, 10 μm. **i,** Quantification of percent area WFA staining. **P < 0.01, *P < 0.05, Two-way ANOVA with post-hoc Bonferroni correction, n=6-13 fields of view (FOVs) per group, from 3 mice per group. Data are presented as mean ± SEM. Complete statistics are provided in Supplementary Table 1.

### Exogenous IL-18 Reduces Stress-Activated Engram Cell Number and Reshapes Their Excitatory Synaptic Input

Memory storage and recall are thought to be supported by a sparse population of activity-recruited neurons, known as engram cells, which are selectively allocated during learning^46^. To investigate whether IL-18 signaling regulates the recruitment and plasticity of stress-activated engram neurons, we combined AAV9-DIO-GFP injection into TRAP2 mice with 4-hydroxytamoxifen (4-OHT)-dependent activity labeling to achieve temporally precise fluorescent tagging of SPS&S-activated neuronal populations (Fig. 5a). Mice received intrahippocampal IL-18 or PBS administration 0.5h before SPS&S exposure and daily after, with 4-OHT delivered 1 h after SPS&S to label the stress-activated ensemble (Fig. 5b). Fifteen days post-SPS&S, obvious GFP fluorescence was detected in the hippocampus of SPS&S-exposed mice compared to non-stressed controls, confirming successful capture of stress-activated neuronal ensembles (Fig. 5c, d and Supplementary Fig. 4a). Strikingly, IL-18 administration markedly reduced the number of GFP-labeled engram cells following SPS&S (Fig. 5c, d), identifying IL-18 as a negative regulator of engram allocation that restricts the competitive recruitment of hippocampal neurons into stress-activated memory ensembles.

**Fig. 5.**
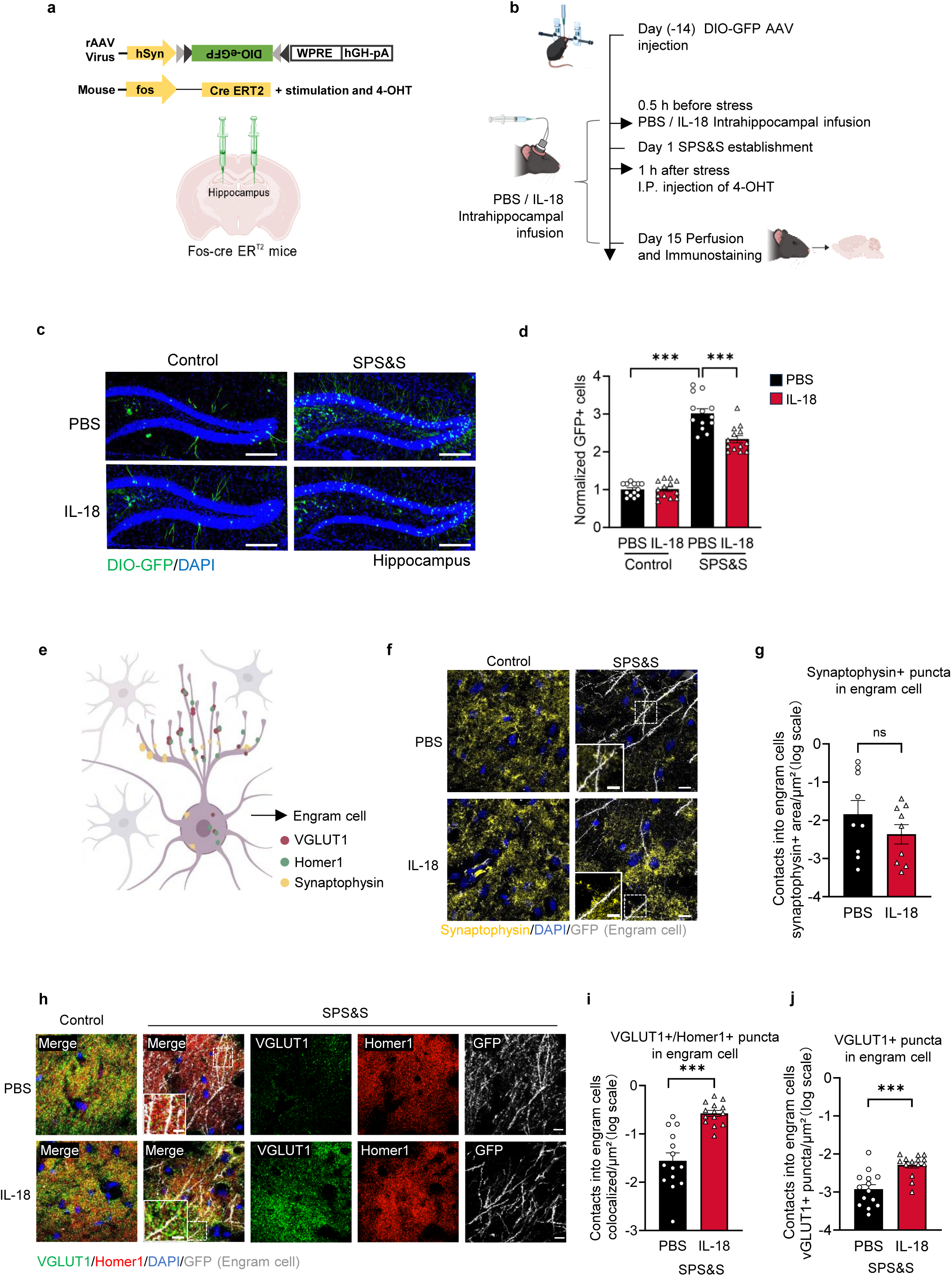
IL-18 modulates fear memory via hippocampal engram cell synaptic remodeling. **a,** Schematic of the TRAP2 engram-labeling system. AAV9-DIO-GFP was locally injected into the hippocampus of Fos-CreER^T2^ mice, enabling activity-dependent fluorescent labeling of SPS&S-activated neurons upon 4-OHT administration. **b,** Experimental timeline. DIO-GFP AAV was injected at day −14; mice received intrahippocampal IL-18 or PBS infusion 0.5h before SPS&S at day 1 and daily after, followed by intraperitoneal 4-OHT injection 1 hour later to complete engram labeling at day1. Brains were collected at day 15 for immunostaining. **c,** Representative confocal images of GFP (green, engram cells) and DAPI (blue) fluorescence in hippocampal sections from control and SPS&S-exposed mice under PBS or IL-18 treatment at day 15. Scale bars, 200 μm. **d,** Quantification of number of eGFP⁺ cells in hippocampus. ***P < 0.001, Two-way ANOVA with post-hoc Bonferroni correction, n=13 fields of view (FOVs) per group, from 3 mice per group. **e,** Schematic illustrating the synaptic markers (VGLUT1, Homer1, synaptophysin) analyzed within GFP⁺ engram cells. **f,** Representative confocal images of synaptophysin (yellow), DAPI (blue), and GFP (engram cell, white) immunostaining in hippocampal engram cells from PBS- and IL-18-treated mice under SPS&S conditions, scale bars, 10 μm. The magnified insets. Scale bars, 5 μm. **g,** Quantification of synaptophysin⁺ puncta contacts in engram cells per unit area (log scale). ns, not significant; unpaired two-tailed t-test; n=9 fields of view (FOVs) per group, from 3 mice per group. The extremely low baseline of engram cells in the control group precluded reliable quantification and thus precluded valid statistical comparisons. **h,** Representative confocal images of VGLUT1 (green), Homer1 (red), DAPI (blue), and GFP (engram cell, white) immunostaining within hippocampal engram cells from PBS- and IL-18-treated under control and SPS&S conditions, scale bars, 10 μm. The magnified insets. Scale bars, 5 μm. **i,** Quantification of VGLUT1⁺/Homer1⁺ co-localized puncta contacts in engram cells per unit area (log scale). ***P < 0.001, Unpaired Student’s t-test, n=14 fields of view (FOVs) per group, from 3 mice per group. **j,** Quantification of VGLUT1⁺ puncta contacts in engram cells per unit area (log scale). ***P < 0.001, Unpaired t-test with Welch’s correction, n=14 fields of view (FOVs) per group, from 3 mice per group. Extremely low baseline engram counts in the control group in panels **i** and **j** preclude reliable quantification (see also panel **g**). Data are presented as mean ± SEM. Complete statistics are provided in Supplementary Table 1.

To examine the effects of IL-18 on engram cell synaptic plasticity following SPS&S, we performed immunofluorescence staining for synaptophysin and VGLUT1/Homer1 within GFP-positive engram cells (Fig. 5e). Synaptophysin puncta density within engram cells was comparable between IL-18-treated and PBS control mice following SPS&S (Fig. 5f, g). In contrast, IL-18 treatment significantly increased VGLUT1 expression and VGLUT1–Homer1 co-localization puncta within engram cell ensembles (Fig. 5h–j), while Homer1 expression alone remained unchanged (Supplementary Fig. 4b), indicating a selective strengthening of presynaptic glutamate release capacity and functional excitatory synaptic connectivity within stress-activated engram cells. Together, these findings demonstrate that IL-18 enhances the excitatory synaptic properties of hippocampal engram cells, extending its broader role in restoring post-traumatic synaptic plasticity to the level of specific memory-encoding neuronal ensembles.

## Discussion

This study reveals an unexpected homeostatic function for IL-18 in the stressed hippocampus, reframing its role in stress-related neuropathology. Canonically regarded as a pro-inflammatory cytokine that amplifies neuroinflammation and exacerbates psychiatric vulnerability, IL-18 is shown here to operate as a neuroimmune regulator that restrains pathological fear memory following traumatic stress. Mechanistically, microglia-derived IL-18 acts on neuronal IL-18R1 to restore stress-impaired synaptic plasticity and remodel perineuronal net architecture, while exogenous IL-18 additionally reshapes the synaptic organization of fear-memory-encoding engram cells. Together, these effects establish a microglia-to-neuron communication axis that opposes maladaptive memory (Supplementary Fig. 4c).

### A context-dependent role for IL-18

The apparent contradiction between IL-18’s canonical pro-inflammatory properties and its protective role here can be reconciled by considering the spatiotemporal context in which signaling operates. Unlike IL-18’s deleterious actions in acute excitotoxic or chronic neurodegenerative settings^19,47^, hippocampal IL-18 under traumatic stress exhibits an induction pattern more consistent with a homeostatic feedback response than with acute inflammatory amplification. A caveat of our design is that IL-18 or IL-18BP administration were initiated before SPS&S and sustained thereafter, spanning both the encoding and consolidation phases of traumatic fear memory and thus precluding formal temporal dissection of its action. Phase-specific interventions will be required to determine whether IL-18 acts primarily during encoding, consolidation, or both^48^.

### Microglia–neuron signaling axis underlying behavioral effects

Cell-type-specific knockdown identified microglia as the principal hippocampal source of IL-18 mediating the behavioral effects in our PTSD model, whereas astrocytic and neuronal IL-18 contributed minimally. Neuronal IL-18R1 was required downstream, defining a microglia-to-neuron signaling axis in which IL-18 released by microglia acts on IL-18R1-expressing neurons. How activation of neuronal IL-18R1 alters synaptic function remains unresolved; candidate mechanisms include modulation of glutamatergic transmission and GABAergic inhibition, which warrants direct investigation in future work.

### Coordinated synaptic and extracellular matrix remodeling

We found that IL-18 signaling reverses stress-induced synaptophysin loss and modulates VGLUT1/Homer1 co-localization, indicating restoration of glutamatergic synaptic integrity. Notably, IL-18 also significantly reduced hippocampal PNN density. PNNs are extracellular matrix structures that stabilize synapses and restrict excessive plasticity, and stress-induced PNN upregulation is thought to stabilize pathological fear memories and confer resistance to extinction. The IL-18-mediated reduction in PNN density may therefore relieve this structural constraint, creating a permissive microenvironment for the synaptic restructuring required to update traumatic associations and facilitate extinction. However, the selective regulation of presynaptic VGLUT1 without a parallel change in postsynaptic Homer1 cannot be fully attributed to PNN remodeling, and points to an additional, PNN-independent effect of IL-18 on presynaptic glutamatergic machinery. Consistent with this interpretation, IL-18R1 knockdown did not further reduce VGLUT1 expression beyond stress-induced levels (Fig. 3g), whereas exogenous IL-18 robustly upregulated VGLUT1 (Fig. 4g), suggesting that endogenous IL-18 signaling under stress is already partially saturated and that supplemental IL-18 drives presynaptic glutamatergic machinery beyond baseline restoration. The molecular pathway by which neuronal IL-18R1 activation regulates VGLUT1 expression remains to be determined. Together, these convergent effects on glutamatergic synapses and the perineuronal matrix identify a dual structural mechanism through which IL-18 may promote neuroplasticity and attenuate the persistence of fear memory.

### IL-18 and hippocampal engram organization

The reduction in GFP-labeled engram cells following IL-18 administration reflects a constraint on engram allocation, the competitive process by which relative intrinsic excitability at the time of encoding determines which neurons are recruited into a memory trace^49,50^. Under normal conditions, this competition maintains engram representations at a characteristic sparseness that supports memory specificity and pattern separation^51^. Traumatic stress disrupts this balance: elevated glucocorticoids persistently increase the excitability and number of activated DG granule cells, expanding engram size in a manner causally linked to fear generalization^52^. The IL-18-mediated reduction in engram cell number may therefore represent a counterbalancing mechanism that opposes stress-induced engram expansion and restores the sparse allocation profile associated with contextually specific memory.

The selective enhancement of glutamatergic connectivity within engram cells may appear counterintuitive, as strengthening synaptic transmission within fear-encoding ensembles might be expected to consolidate rather than attenuate fear memory. However, studies of hypothalamic oxytocin engram neurons demonstrate that a fear-experience-dependent increase in glutamate release within a specific engram population is both sufficient and necessary for attenuating contextual fear^53^, establishing that enhanced glutamatergic output within engram cells can serve a fear-suppressing function. Together, IL-18 influences engram configuration at two levels: limiting the number of neurons recruited during traumatic stress encoding and selectively strengthening glutamatergic connectivity within those that are allocated. Whether these two effects arise from a common mechanism or represent parallel actions of IL-18 signaling on distinct cellular targets remains an open question for future.

### Translational considerations

Translating these findings into therapy will require caution. In chronic neuroinflammation or different brain regions (such as the amygdala), IL-18 may exert pathogenic effects through alternative mechanisms^37^. This complexity demands that future therapeutic strategies must be precisely targeted (e.g., specific cellular sources, specific receptor subtypes, specific brain regions). Our research suggests that enhancing IL-18 signaling within the hippocampus or mimicking its downstream neuroprotective effects may represent a novel direction for PTSD treatment. However, considering IL-18’s bidirectional nature, administration timing, dosage, and delivery methods require strict control.

In summary, microglial IL-18 is not only an alarm signal; but it limits pathological fear memory by reshaping synaptic integrity, perineuronal net architecture, and the size of stress-activated engram ensembles, prompting a reconsideration of how the brain mobilizes its own immune machinery after trauma. The microglia-to-neuron IL-18 axis exemplifies how neuroimmune communication can act not as a peripheral modulator of brain function but as an integral organizer of adaptive responses to stress, and may inform therapeutic strategies for other stress-related disorders.

## Acknowledgments

We thank Professor Tao Li for help discussion.

## Author Contributions

W.H.L., T.L., and X.Y.F. designed the study. X.Y.F. performed all experiments with guidance from T.L. and W.H.L. X.Y.F. established the SPS&S model with technical assistance from G.B.X. and X.H. H.C., Y.N.W., R.L., and Y.Z.S. assisted with image quantification and statistical analysis. Q.Y.H., X.X., K.W., L.B.Y. and J.W.D. provided technical assistance with behavioral testing and confocal imaging. T.Z. and A.L.L. contributed to interpreting the results. T.L. and W.H.L. supervised the study, provided conceptual guidance. T.L., X.Y.F., X.H., and W.H.L. wrote the manuscript. All authors reviewed and approved the final manuscript.

## Methods

### Animals

All experiments used 8–12-week-old male and female C57BL/6 mice. Animals were housed under specific pathogen-free (SPF) conditions with a 12-h light/dark cycle (lights on 08:00–20:00, 50 lux), at 22°C and 40–60% relative humidity, with ad libitum access to food and water unless otherwise stated. For IL-18 reporter experiments, IL-18-Luc-2A-EGFP reporter mice were used to visualize IL-18-expressing cells via GFP immunostaining and luciferase activity (Shanghai Model Organisms Center). For engram-labeling experiments, Fos2A-iCreER (TRAP2, JAX:030323) mice were obtained from Jackson Laboratory. Adult C57BL/6 mice were obtained from Beijing Vital River Laboratory Animal Technology. All procedures were approved by the Institutional Animal Care and Use Committee of Nanhu Laboratory and conducted in accordance with institutional guidelines.

### Drugs and reagents

Recombinant murine IL-18 (Med Chem Express, MCE) was dissolved in sterile phosphate-buffered saline (PBS) and stored at −80°C until use. IL-18 binding protein (IL-18BP; MCE) was dissolved in sterile PBS and stored at −80°C. 4-Hydroxytamoxifen (4-OHT; MCE) was dissolved in corn oil to a final concentration of 10 mg/mL immediately before use. All drug solutions were prepared fresh or aliquoted to avoid repeated freeze–thaw cycles.

### Plasmid and virus generation

AAV vectors were generated by standard cloning methods and packaged by the indicated manufacturers. For cell-type-specific IL-18 knockdown, AAV-Iba1-shIL18, AAV-GFAP-shIL18, AAV-hSyn-shIL18, and AAV-CaMKII-shIL18 were constructed by cloning validated shRNA sequences targeting mouse Il18 into AAV backbone vectors containing cell-type-specific promoters (shIL-18: 5′-AGAGTCTTCTGACATGGCAGC-3′). Scramble shRNA (EF1-shCon) served as a control (5′-CCTAAGGTTAAGTCGCCCTCG-3′, BrainVTA). Corresponding constructs targeting Il18r1 (AAV-Iba1-shIL18R1, AAV-GFAP-shIL18R1, AAV-hSyn-shIL18R1) were generated using validated shRNA sequences (shIL-18R1: 5′-TGTGAAGCACTGCCTTTGAAC-3′, BrainVTA). For engram-labeling experiments, AAV2/9-hSyn-DIO-EGFP-WPRE-hGH polyA (AAV9-DIO-GFP, 5.28 × 10¹² gc/mL, BrainVTA) was used to label Cre-expressing (activity-activated) neurons in TRAP2 mice. All viruses were diluted to 3–8 × 10^12^ gc/mL in sterile PBS prior to injection.

### Stereotaxic surgery

Mice were anesthetized with isoflurane and positioned in a stereotaxic frame (RWD Life Science). Coordinates were identified using an online mouse brain atlas (https://labs.gaidi.ca/mouse-brain-atlas). For bilateral hippocampal AAV delivery, viruses were injected into the dorsal hippocampus (AP −2.2, ML ±1.3, DV −2 mm; 250 nL per hemisphere at 50 nL/min). Under stereotaxic anesthesia, bilateral guide cannulae (RWD Life Science) were implanted into the dorsal hippocampus (AP −2.2, ML ±1.3, DV −1.6 mm). Mice were allowed to recover for at least 21 days (shRNA AAV injection) or 14 days (cannula implantation) before experimental procedures.

### Intrahippocampal infusion

Bilateral intrahippocampal infusions were performed via implanted cannulae using an RWD microinfusion pump. IL-18 (50 ng/side), IL-18BP (50 ng/side), or vehicle (PBS) was delivered in 0.25 μL at a rate of 0.1 μL/min, with cannulae left in place for an additional 10 min to ensure complete diffusion. All infusions were administered 30 min prior to SPS&S exposure on the day of treatment, or 30 min before behavioral testing on subsequent days. IL-18 was administered daily from the day of SPS&S through day 15. To ensure adequate suppression of endogenous IL-18 prior to traumatic stress exposure, IL-18BP infusions began 3 days before SPS&S and continued daily through day 15.

### 4-OHT administration for engram labeling

To label SPS&S-activated hippocampal neurons in TRAP2 mice, 4-OHT (75 mg/kg) was administered intraperitoneally 1 hour after completion of the SPS&S procedure. This timing ensures that CreER^T2^ nuclear translocation occurs specifically within the window of stress-activated neuronal activity, enabling temporally precise labeling of engram-competent neurons.

### Behavioral assays

#### Single prolonged stress and shock (SPS&S) model

The SPS&S paradigm was established according to the protocol described by Xi et al. with some modification^1^. Briefly, mice were subjected to four consecutive stressors in a single session: (1) 2-hour restraint in a well-ventilated restraint tube; (2) 15-minute forced swim in a cylinder (diameter 15 cm, depth 25 cm) filled with water at 23–25°C; (3) brief diethyl ether anesthesia until loss of righting reflex; and (4) contextual foot shock (0.8 mA, constant current, 10 s) delivered after 5 min placement into the shock chamber. Following SPS&S, mice were returned to their home cages and remained undisturbed until behavioral testing at days 8 and 15 post-stress. Control mice were handled identically but received no stressors.

#### Open field test

To assess locomotor activity behavior, mice were placed in a square open-field arena (50 × 50 cm) and allowed to explore freely for 10 min. Total distance traveled was recorded using LabState Software (AniLab)

#### Elevated plus maze (EPM)

The EPM apparatus consisted of two open arms (25 × 5 cm, 0.5-cm raised edges) and two enclosed arms (25 × 5 × 16 cm) arranged in a plus configuration, elevated 50 cm above the floor with a central platform (5 × 5 cm). Mice were placed in the center facing an open arm and allowed to explore freely for 10 min. Time spent in open arms and number of open-arm entries were recorded as indices of anxiety. The maze was cleaned with 70% ethanol between animals. Behavior was tracked using SMART Video Tracking System (Panlab).

#### Tail suspension test (TST)

After a 1-h habituation period in the test room, mice were individually suspended by their tails using adhesive tape attached to a bar 50 cm above the floor in a tail-suspension box (20 × 20 × 32 cm) for 10 min. Immobility time during the last 8 min was recorded using SMART Video Tracking System (Panlab) as an index of depressive-like behavior.

#### IL-18 luciferase reporter assay

To quantify region-specific IL-18 bioactivity, hippocampal, medial prefrontal cortex (mPFC), and basolateral amygdala (BLA) tissues were dissected from IL-18-Luc-2A-EGFP reporter mice on day 3 post-SPS&S or from controls. Tissue was homogenized in passive lysis buffer (Promega) and centrifuged at 12,000 rpm for 10 min at 4°C. Supernatants were transferred to white-bottom 96-well plates and luciferase activity was measured using the Luciferase Assay System (Promega) on a luminometer (SpectraMax, Molecular Devices). Corrected luciferase activity was normalized to total protein concentration determined by Bradford assay (Thermo Fisher Scientific).

#### ELISA

Dorsal hippocampal tissues were dissected and flash-frozen at multiple time points post-SPS&S (0, 2, 4, 8, 24 h, 3, 8, 15 days). Tissues were homogenized in T-PER tissue protein extraction reagent (Thermo Fisher Scientific, 78510) supplemented with protease and phosphatase inhibitor cocktails (cOmplete and PhosSTOP, Roche). Lysates were centrifuged at 15,000 rpm for 10 min at 4°C, and supernatants were collected. Active (mature) IL-18 and IL-1β protein concentrations were measured using commercially available ELISA kits specific for the cleaved form of IL-18 and IL-1β(mouse IL-18 ELISA kit, MBL; mouse IL-1β ELISA kit, Thermo Fisher Scientific) according to the manufacturers’ instructions. All values were normalized to total protein concentration.

#### Quantitative RT-PCR (qPCR)

Total RNA was extracted from hippocampal tissue using TRIzol reagent (Sigma, T9424) following the manufacturer’s protocol. Total RNAs (500 ng) were reverse-transcribed to cDNA using the PrimeScript RT Reagent Kit (RR037A, Takara). mRNA expression was analyzed by qPCR with PowerUp SYBR Green Master Mix (A25742, Thermo Fisher Scientific) on an Applied Biosystems StepOnePlus system. The primer sequences used for qRT–PCR were as follows: IL-18, forward 5′-GACAGCCTGTGTTCGAGGAT-3′ and reverse 5′-TGGATCCATTTCCTCAAAGG-3′; Actb served as the reference gene, forward 5′-CATTGCTGACAGGATGCAGAAGG-3′ and reverse 5′-TGCTGGAAGGTGGACAGTGAGG-3′.

#### Immunofluorescence

Mice were transcardially perfused with ice-cold PBS followed by 4% paraformaldehyde (PFA) in PBS. Brains were post-fixed in 4% PFA overnight at 4°C, cryoprotected in 30% sucrose in PBS until equilibrated (≥48 h), and cryosectioned at 30 μm on a Leica cryostat. For immunostaining, air-dried frozen sections mounted onto glass slides were blocked for 1 h at room temperature in blocking buffer (3% goat serum, 1% BSA, 0.3% Triton X-100 in PBS), and incubated overnight at 4°C with primary antibodies diluted in blocking buffer. Primary antibodies used in this study included: chicken anti-NeuN (1:1000, Millipore, ABN91), rabbit anti-Iba1 (1:1000, FUJIFILM Wako, 019-19741), mouse anti-GFAP (1:500, Thermo Fisher Scientific, 130300), rabbit anti-synaptophysin (1:200, Cell Signaling Technology, 36406T), goat anti-mouse IL-18 receptor α (1:100, R&D Systems, AF856), rat anti-mouse IL-18 receptor α/IL-1 R5 (1:100, R&D Systems, MAB12161-100), biotinylated Wisteria floribunda agglutinin (WFA) (1:500, Sigma-Aldrich, L1516), rabbit anti-S100β (1:500, Proteintech, 15146-1-AP), rabbit anti-Homer1 (1:500, Synaptic Systems, 160003), rabbit anti-VGLUT1 (1:1000, Synaptic Systems, 135511), rabbit anti-aggrecan (1:500, Millipore, AB1031), NeuroTrace 640/660 deep-red fluorescent Nissl stain (1:500, Thermo Fisher Scientific, N21483), and chicken anti-GFP (1:1000, Aves Labs, GFP-1020). Sections were washed three times (10 min each) with 0.05% Tween-20 in PBS, then incubated with matched Alexa Fluor-conjugated secondary antibodies (488, 546, 647; Thermo Fisher; 1:1000) for 1 h at room temperature. For WFA, streptavidin-conjugated Alexa Fluor 647 was used. Sections were washed, counterstained with DAPI (1:1000, Millipore Sigma), mounted with ProLong Gold anti-fade mountant (Thermo Fisher, P36930), and stored at 4°C in the dark. Images were acquired on Olympus VS200 and Zeiss 900 confocal microscope.

#### Image analysis and quantification

For synaptophysin, VGLUT1, Homer1, and WFA quantification, confocal z-stack images (8-10 optical sections, 0.5-μm step) were acquired from the hippocampal CA1 region. Puncta density was quantified using the ’Analyze Particles’ function after a uniform threshold in the same group (ImageJ, NIH). Co-localization of VGLUT1 and Homer1 puncta was assessed using the ImageJ Colocalization plugin, with co-localized puncta defined as overlapping signals within ≤0.5 μm. WFA staining intensity was expressed as percent area above threshold. For engram cell analyses, GFP^+^ cells were identified by manual counting across 3–5 hippocampal sections per mouse. Synaptophysin^+^, VGLUT1^+^, and Homer1^+^ puncta contacts in GFP^+^ engram cell somata and proximal dendrites were quantified and normalized to engram cell area (μm^2^; expressed on a log scale). For IL-18-GFP cell analyses in whole brain, images were analyzed using Imaris (9.8.0) software.

### Statistical analysis

Statistical analyses were performed using GraphPad Prism 10.1.2 (GraphPad Software). Data normality was assessed using the D’Agostino–Pearson or Shapiro–Wilk tests. For two-group comparisons, unpaired or paired Student’s *t*-tests were applied to normally distributed data; the Mann–Whitney U or Wilcoxon signed-rank tests were used for non-normal distributions. Multiple-group comparisons used one-way ANOVA with Bonferroni post hoc correction (normal distribution) or Kruskal–Wallis test with Dunn’s post hoc correction (non-normal distribution). A two-way ANOVA was performed, followed by post hoc tests with Bonferroni or Holm-Šídák correction as appropriate. All data are presented as mean ± SEM. Statistical significance thresholds: \**P* < 0.05, \*\**P* < 0.01, \*\*\**P* < 0.001; ns, not significant. Sample sizes (*n*) indicate the number of animals per group and are reported in the figure legends. Mice were randomly assigned to treatment groups, and investigators were blinded to genotype and treatment conditions during behavioral testing and image analysis. All key experiments were reproduced in at least two independent cohorts.

---

**Supplementary Fig. 1.**
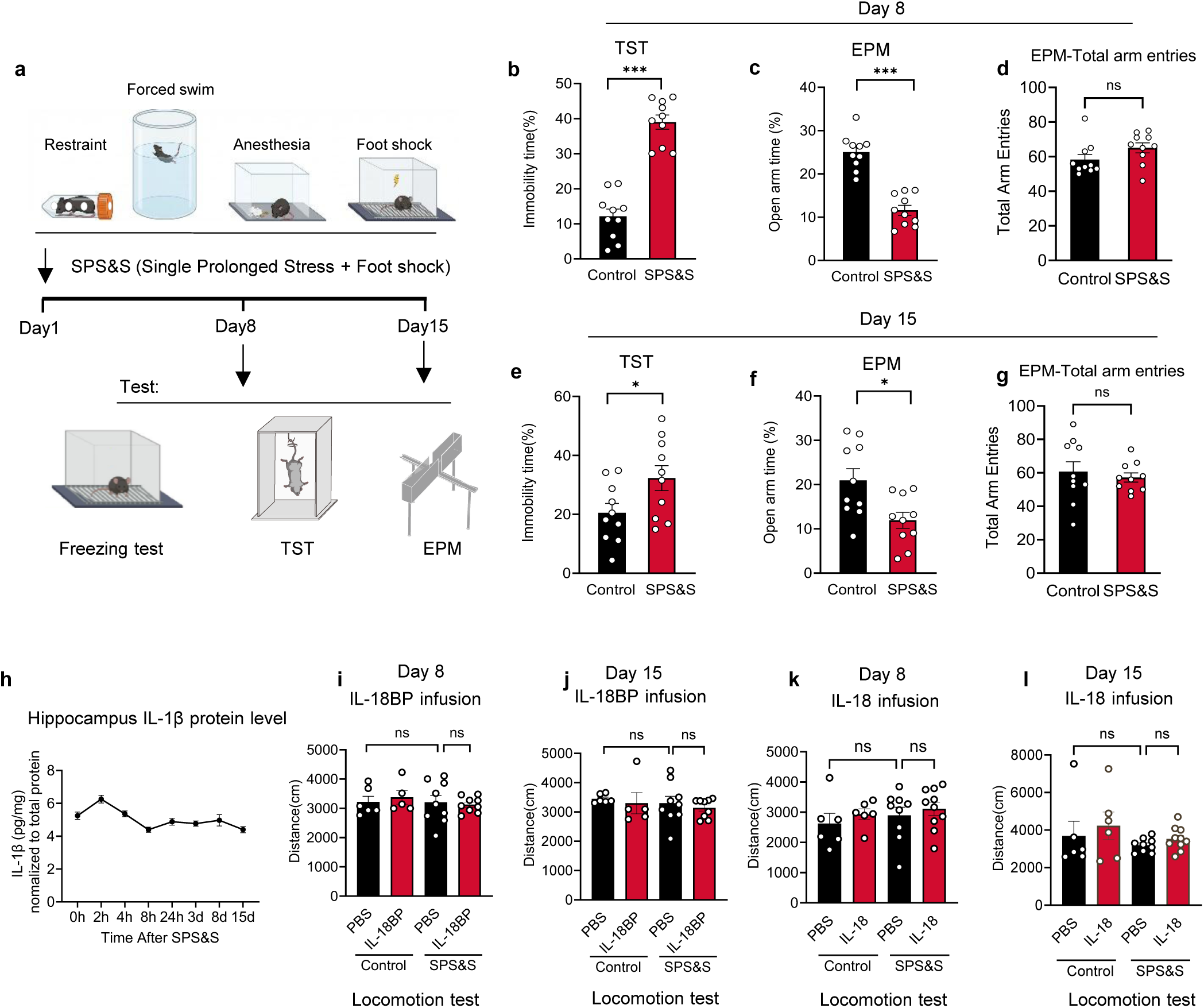
SPS&S induces PTSD-like behavioral and neuroimmune changes, and IL-18 pathway modulation does not alter locomotor activity. **a,** Schematic of the SPS&S modeling procedure (restraint, forced swim, anesthesia, foot shock) and behavioral testing timeline, including contextual fear freezing test, tail suspension test (TST), and elevated plus maze (EPM) at days 8 and 15 post-stress. **b,** TST immobility time at day 8 in control and SPS&S mice. ***P < 0.001; unpaired two-tailed t-test; n = 10 mice per group. **c,** EPM open arm time at day 8 in control and SPS&S mice. ***P < 0.001; unpaired two-tailed t-test; n = 10 mice per group. **d**, EPM Entries: ns, not significant, Mann–Whitney U test, n=10 mice per group; **e**, TST immobility time at day 15 in control and SPS&S mice. *P < 0.05; unpaired two-tailed t-test; n = 10 mice per group, **f**, EPM open arm time at day 15 in control and SPS&S mice. *P < 0.05; unpaired two-tailed t-test; n =10 mice per group. **g**, EPM Entries, ns, not significant, Unpaired t-test with Welch’s correction. n=10 mice per group. **h**, Temporal profile of hippocampal IL-1β protein levels (normalized to total protein) from 0 to 15 days post-SPS&S. One-way ANOVA with Dunnett’s post hoc test; n = 7-11 per time point. **i**, Total locomotor distance at day 8 in mice receiving intrahippocampal IL-18BP or PBS following SPS&S or NT conditions, ns, not significant Two-way ANOVA with post-hoc Bonferroni correction, n=5-9 mice per group. **j**, Same as i at day 15, ns, not significant, Two-way ANOVA with post-hoc Bonferroni correction, n=5-9 mice per group. **k**, Total locomotor distance at day 8 in mice receiving intrahippocampal IL-18 or PBS following SPS&S or NT conditions, ns, not significant, Two-way ANOVA with post-hoc Bonferroni correction, n=6-10 mice per group. **l**, Same as k at day 15, ns, not significant, Two-way ANOVA with post-hoc Bonferroni correction, n=6-10 mice group. Data are presented as mean ± SEM.

**Supplementary Fig. 2.**
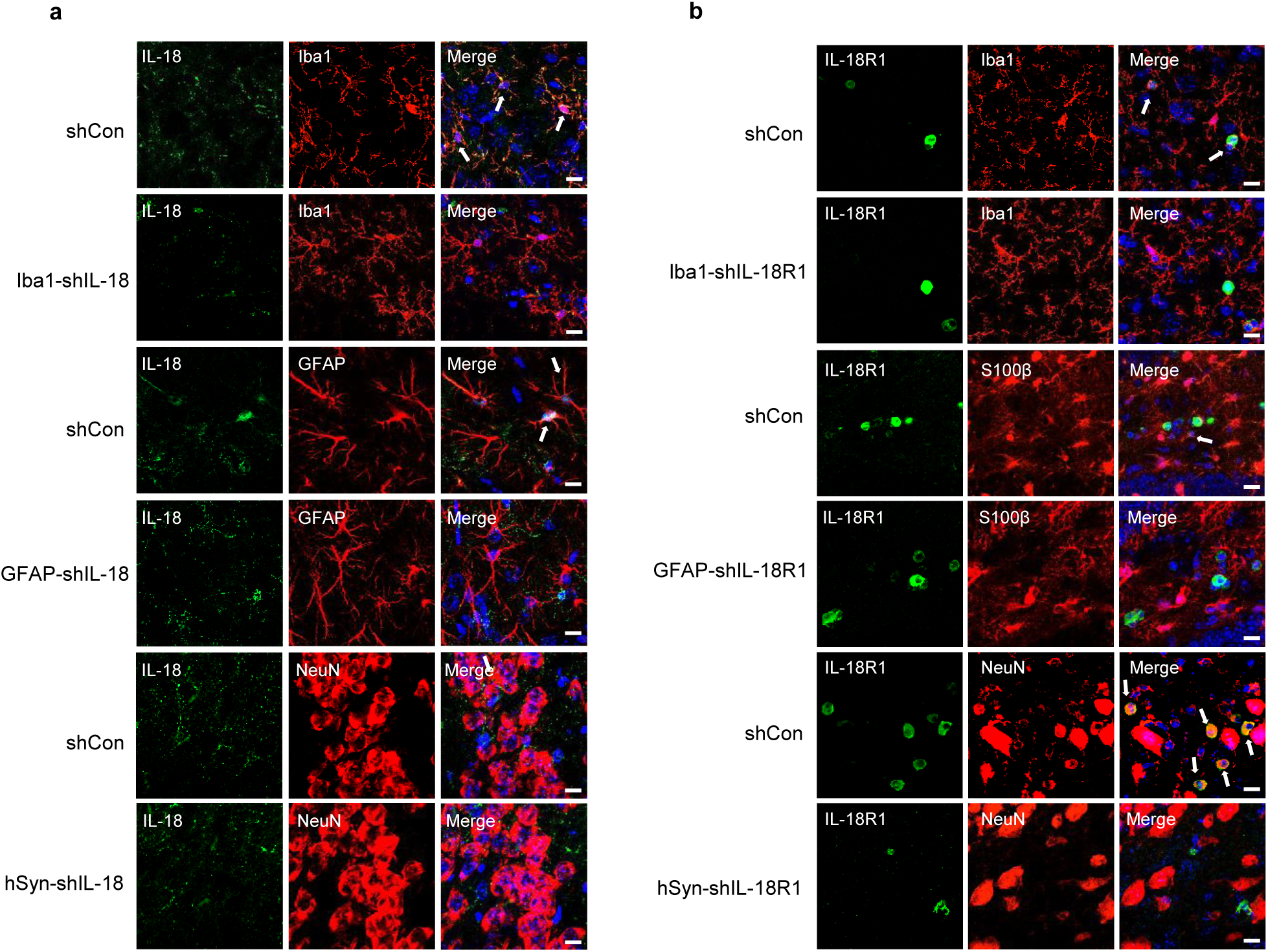
Cell-type-specific validation of IL-18 and IL-18R1 knockdown efficiency in the hippocampus. **a,** Representative confocal images validating cell-type-specific AAV-mediated IL-18 knockdown. Left panels show co-immunostaining for IL-18 (green) with Iba1 (red, microglia) or GFAP (red, astrocytes) or NeuN (red, neurons) in shCon, Iba1-shIL18, GFAP-shIL18, and hSyn-shIL18 groups. Merged images confirm selective depletion of IL-18 in the targeted cell type. White arrows indicate co-localization. Scale bars, 10 μm. **b,** Representative confocal images validating cell-type-specific AAV-mediated IL-18R1 knockdown. Right panels show co-immunostaining for IL-18R1 (green) with Iba1 (red), S100β (red), or NeuN (red) in shCon, Iba1-shIL18R1, GFAP-shIL18R1, and hSyn-shIL18R1 groups. White arrows indicate residual or depleted co-localization in the targeted cell type. Scale bars, 10 μm.

**Supplementary Fig. 3.**
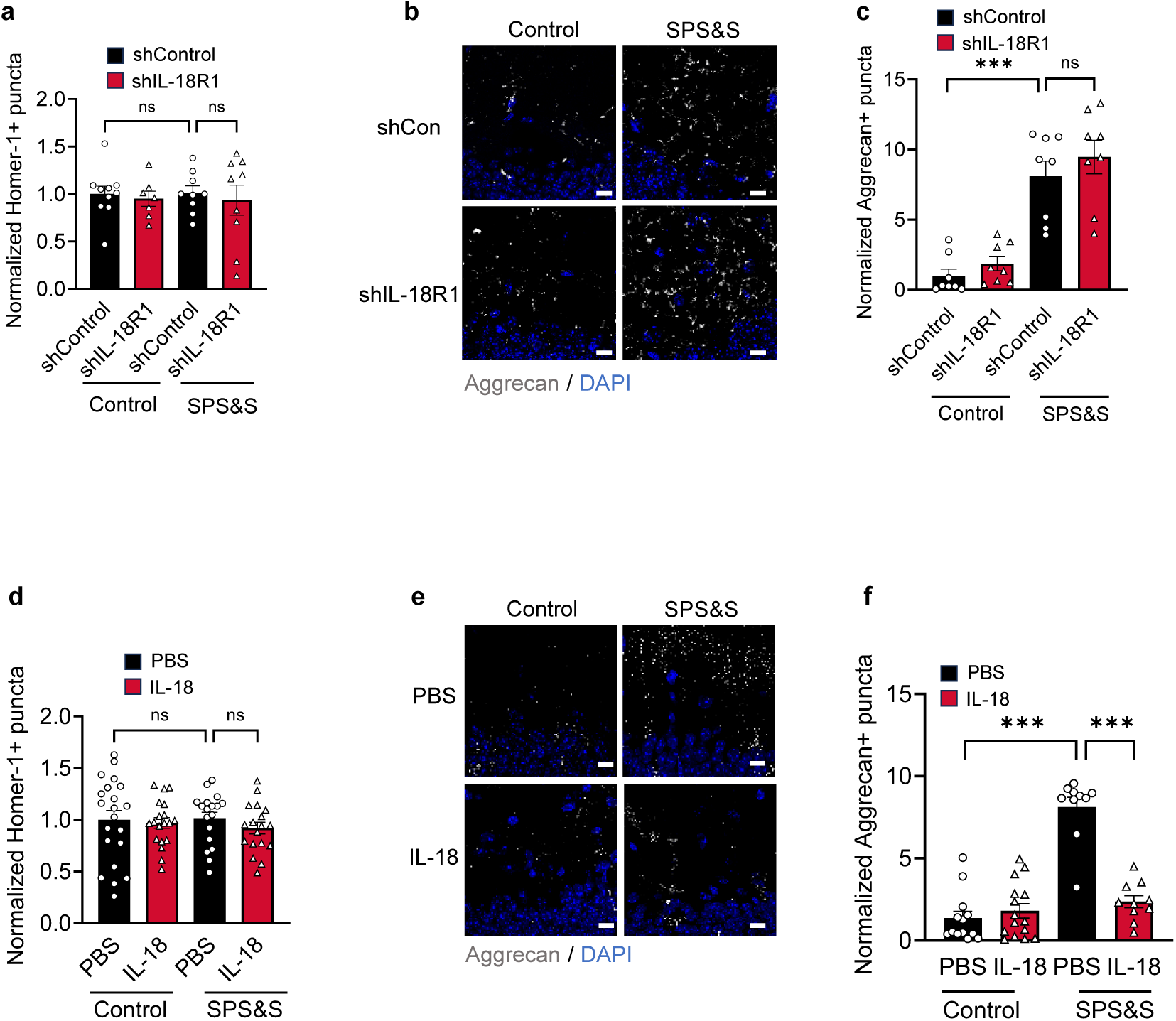
Homer1 expression is unaffected by IL-18R1 knockdown or IL-18 treatment. **a,** Quantification of normalized Homer1⁺ puncta density in hippocampal CA1 from shControl and shIL-18R1 mice under control and SPS&S conditions. ns, not significant; Two-way ANOVA with post-hoc Holm-Šídák, n=7-10 fields of view (FOVs) per group, from 3 mice per group **b,** Representative confocal images of aggrecan (white) and DAPI (blue) immunostaining in hippocampal CA1 from shControl and shIL-18R1 mice under control and SPS&S conditions. Scale bars, 10 μm. **c,** Quantification of normalized aggrecan⁺ puncta density. ***P < 0.001, ns, not significant, Two-way ANOVA with post-hoc Holm-Šídák, n=8 fields of view (FOVs) per group, from 3 mice per group. **d,** Quantification of normalized Homer1⁺ puncta density in hippocampal CA1 from PBS- and IL-18-treated mice under control and SPS&S conditions. ns, not significant; two-way ANOVA with post-hoc Holm-Šídák, n=18-21 fields of view (FOVs) per group, from 3 mice per group **e,** Representative confocal images of aggrecan (white) and DAPI (blue) immunostaining in hippocampal CA1 from PBS-and IL-18-treated mice under control and SPS&S conditions. Scale bars, 10 μm. **f,** Quantification of normalized aggrecan⁺ puncta density. ***P < 0.001, Two-way ANOVA with post-hoc Holm-Šídák, n=10-15 fields of view (FOVs) per group, from 3 mice per group. Data are presented as mean ± SEM.

**Supplementary Fig. 4.**
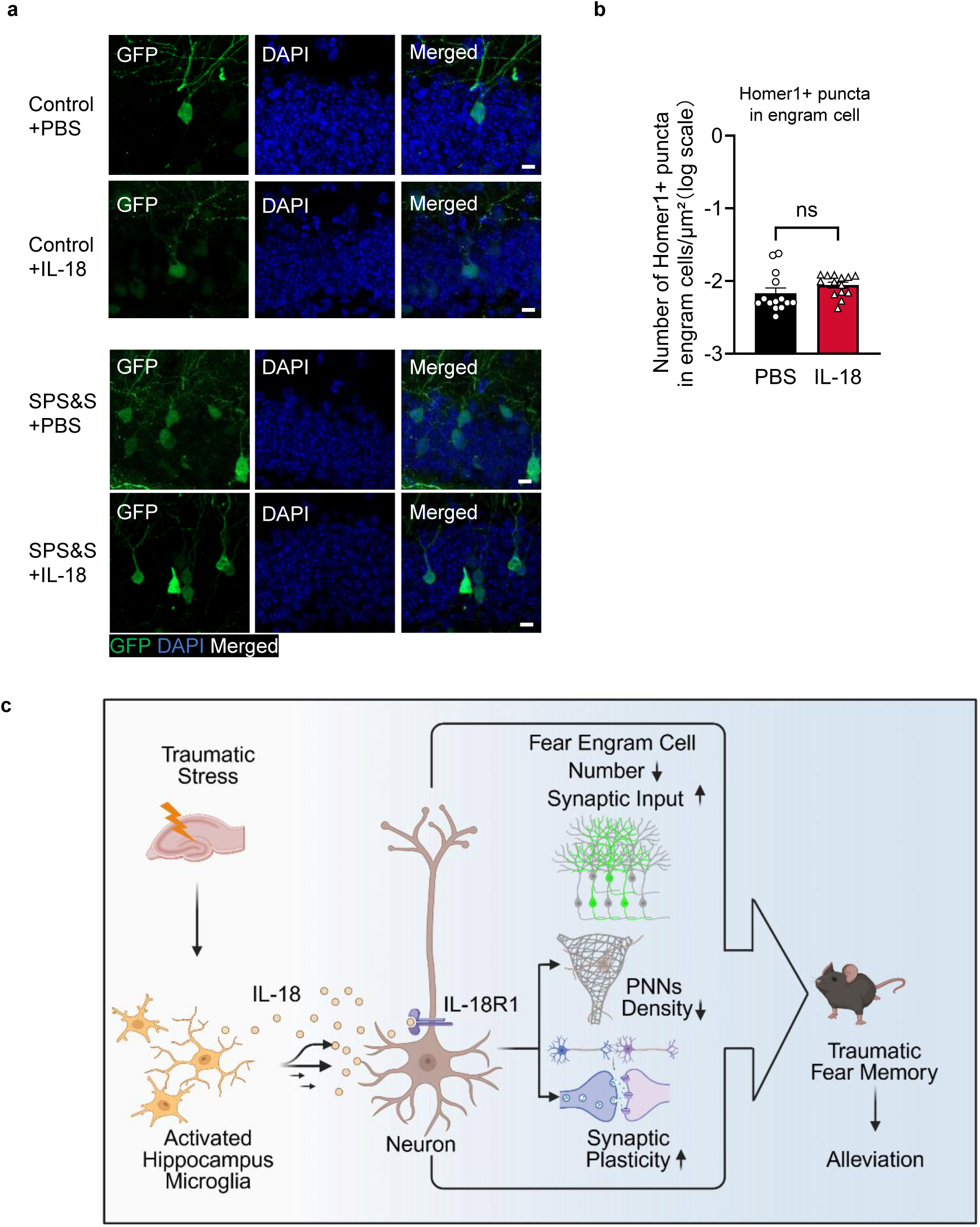
Engram cell labeling validation and Homer1 expression in hippocampal engram cells following IL-18 treatment. **a,** Representative confocal images of GFP (green) and DAPI (blue) fluorescence in hippocampal sections from PBS- and SPS&S-exposed TRAP2 mice at day 15, confirming successful activity-dependent engram cell labeling in the dentate gyrus with axonal projections to CA1. Scale bars, 10 μm. **b,** Quantification of Homer1⁺ puncta contacts in GFP⁺ engram cells per unit area (log scale) in PBS- and IL-18-treated SPS&S mice. ns, not significant; Mann–Whitney U test, n=14 fields of view (FOVs) per group, from 3 mice per group. **c,** Schematic summarizing the microglia-to-neuron IL-18 signaling axis and its therapeutic effects on PTSD-associated circuit impairment. Following traumatic stress, microglial-derived IL-18 signals through neuronal IL-18R1 to restore impaired synaptic plasticity, reduce perineuronal net (PNN) density in the extracellular matrix, and modulate hippocampal engram ensemble synaptic organization, collectively constraining maladaptive fear memory consolidation.

